# A summary of intraspecific size variation for large odontocetes

**DOI:** 10.1101/2024.09.05.610699

**Authors:** Joseph McClure

## Abstract

The study of cetacean size and growth aids in research about their energetics, ecology, evolution, and taxonomy. Data on total body length and mass of large cetaceans is retained in an overwhelming amount of literature and previous reviews are limited in discussing the variation of both length and mass. Nearly all the known published literature was examined to provide a brief but comprehensive summary of the adult size of four large odontocetes. To supplement the existing literature, data from the International Whaling Commission catch database were reviewed and incorporated into analyses to improve descriptions of intraspecific variation in adult size for *Physeter macrocephalus*. Regional comparisons of commercial data for *P. macrocephalus* revealed apparent differences in adult size that are likely more attributable to catch biases rather than a diagnostic difference in size between populations.

## 1 Introduction

Cetaceans represent some of the largest known animals (Lockyer, 1976). Body size and growth is a prominent biological character in studies on the cetacean ecology (Pershing et al., 2010), conservation (Christiansen et al., 2020), stock divisions (Fung, 2016; Ichihara, 1966; Kasuya & Tai, 1993; Pastene et al., 2020), and evolution (Bianucci et al., 2019; Bisconti et al., 2021). Popular megafauna are often prone to misinformation regarding both their typical and maximum confirmed body size (De Maddalena et al., 2003; Matthews & Parker, 1950; Nielsen, 2017; Paxton, 2016; Roberts, 2012; Romanov et al., 2018). Searching and compiling size data is cumbersome with some works being nearly lost in obscurity, particularly literature with limited foreign translations.

Total body length (TBL) and mass are the two most important and widely-used measurements for body size in cetaceans and quantifying their variation can provide insight about the environmental influences on a species, as it has for birds and terrestrial mammals (Plavcan, 2012; Zheng et al., 2023). The maximum attainable size, as a limit of adult size variation, is also an important parameter for drawing inferences on survival, longevity, and growth (Trites & Pauly, 1998).

Immature and mature individuals were not separated in the datasets of previous reviews of cetacean body size variation (MacLeod, 2006; McClain et al., 2015), making their results less reliable for research that typically isolates adults to remove ontogenetic influences (Bartle & Sagar, 1987; Brecko et al., 2008). Furthermore, the investigation of body mass data was minimal and is particularly under-studied in ziphiids. This is particularly concerning as mass is an important variable for calculating the energetics of feeding, growth, and reproduction (Christiansen et al., 2022; Lockyer, 1981a, 1981b; New et al., 2013).

It was therefore a great priority for the author to comprehensively review the body size for four of the largest odontocete species: *Orcinus orca* (Linnaeus, 1758), *Berardius bairdii* (Stejneger, 1883), *Hyperoodon ampullatus* (Forster, 1770), and *Physeter macrocephalus* (Linnaeus, 1758). Each species appears to wean and initiate reproductive development at similar ages (Benjaminsen & Christensen, 1979; Feyrer et al., 2020; Kasuya, 1997; Newsome et al., 2009; Ohsumi, 1965; Olesiuk et al., 2005). However, *O. orca* and ziphiids physically mature at 15-20 years (Benjaminsen & Christensen, 1979; Christensen, 1984; Kasuya, 1997), while physical growth for *P. macrocephalus* ceases after 30 and 50 years in females and males respectively (Clarke et al., 1994, 2011). Comparisons among these species will aim to examine if the interval between sexual and physical maturity is correlated with the relative variation in adult size.

Both literature and the data within the International Whaling Commission (IWC) catch database (CAllison, 2020) were analyzed to obtain adult length distributions and body mass relationships. These species were among the most extensively researched throughout commercial whaling and their large size and legal protections limit the collection of additional morphometric data. This condensing of over a century of information may also prove convenient for researchers developing ecological, energetic, or phylogenetic models that benefit from incorporating intraspecific information about TBL and body mass.

## 2 Methods

Nearly all accessible literature on biological examinations, strandings, museum specimens, and whaling data were surveyed solely by the author via mainstream journals, prominent gray literature available online (Reports of the IWC, Institute of Cetacean Research, Discovery Committee, etc.), reference libraries, purchases of antiquarian books, and personal communications from researchers to uncover information on postnatal growth, length, and body mass. Primary search terms were ‘total length’, ‘’photogrammetry’’, ‘growth curve’, ‘age data’, ‘maximum girth’, ‘weight/mass’, and ‘oil yields’. This review of literature aims to provide parameters on adult size, emphasizing the average length at sexual maturity (*L_sm_*), average asymptotic size (*L*_∞_) and maximum size (*L*_max_) for the species being reviewed. Data on the size at birth (*L*_b_) and weaning (*L*_w_) are also provided as relevant parameters influenced by adult female size.

Literature containing morphometric data or growth parameters were selected on the basis of four criteria. First, all available primary literature on individual morphometric data (TBL, girth, individual oil yields, body mass, and body mass formulas) were prioritized to ensure accuracy. All redundant secondary sources that provided no revisions or insights to existing data were excluded. Secondly, TBLs reported in literature had to correspond to the standard length, which is from the tip of the upper jaw to notch of the flukes along a straight line. This was assumed for all measurements reported by scientists during the timeframe the convention had been widely adopted (ca. 1920). Data from older literature were excluded unless the explicit procedure of measuring or other material were reported to determine the reliability or allow the conversion of measurements. Thirdly, literature where all body mass data were either explicitly stated in the text or otherwise determined by the current author to be purely estimates rather than those measured more directly (piecemeal, volumetric, or intact weighings) were excluded. Lastly, for species or populations with multiple existing estimates for growth parameters, only literature providing the most updated or precisely calculated parameters was selected. In total, 323 sources from literature (69 books, 6 conference papers, 229 articles published in journals and grey literature, 10 technical reports/research documents, 6 theses, and 3 media articles) were surveyed and 153 were selected after filtering in accordance with the above criteria.

Throughout the review, body mass formulas for whales were developed by the least-squares fit of the log-transformed values into a linear regression (Equation 1), which is then converted to the power function presented in Equation 2 (Schultz, 1938). Most of the published body mass data for larger whales were the sums of individual pieces weighed on platform scales or bulk fillings of pressure cookers (Lockyer, 1976). Due to fluid loss and minor losses of solid tissue, piecemeal weighings underestimate the true body mass. A correction factor of at least 12% was empirically determined for *P. macrocephalus* from comparative weighings of intact and flensed carcasses (Gambell, 1970; Lockyer, 1991)

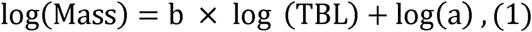

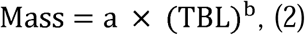

This review also provides the Rice-Wolman (RW) mass prediction model (Equation 3) that incorporates girth (Rice & Wolman, 1971). The inclusion of girth as an additional variable enables the RW model to more precisely estimate the body mass of a whale (George, 2009; George et al., 1990; Rice & Wolman, 1971) even with small samples sizes (*n* < 10). When the raw data are available, comparisons between body mass relationships are made by using Akaike information criterion (AIC) model selection (Burnham & Anderson, 2002). Body mass data, references, and code for model selection are contained in Supplementary Files 2-4, respectively.

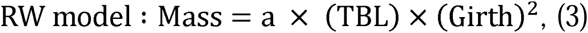

In this review, mean adult length (*L*_mean_) is described as the average length of the sexually mature population, which is distinct from the *L*_∞_ parameter. Since large cetaceans are sexually mature well before physical growth ceases (Bryden, 1972), the *L*_mean_ should conceptually be slightly lower than the *L*_∞_, as observed in biological examinations for baleen whales (Ohsumi et al., 1958; Zenitani et al., 2001). The analyses for *P. macrocephalus* within the IWC database are restricted to catches from after the introduction of minimum length regulations in 1937, corresponding to the ‘’Late period’’ as described in previous work on southern blue whales (Branch et al., 2007). This convention helps minimize data quality issues within early whaling data such as non-standard measuring techniques, visual estimation, and rounding to 5-ft intervals (Branch et al., 2007; Donovan, 2000).

With the exception of the revised data for Soviet pelagic catches in the Southern Hemisphere (Yablokov, 1994; Zemsky et al., 1995), all data for *P. macrocephalus* reported from Soviet and Japanese operations prior to 1972 were filtered out due to the widespread data falsification prior to the enforcement of the International Observer’s Scheme (IOS) (Clapham & Ivashchenko, 2016; Ivashchenko et al., 2014; Ivashchenko & Clapham, 2015). The intentional ‘stretching’ of whales below the minimum legal size by other nations is also known to have affected the female distributions for *P. macrocephalus* prior to the IOS (Best, 1989). To accommodate for the possible influence of these artefacts, catches taken before and after 1972 were analyzed separately.

Biological examinations and age-length keys from the commercial catches indicate that the length distributions for mature whales should be approximately normal (Branch et al., 2007; Ohsumi, 1977; Ohsumi et al., 1958). This can be directly observed for the females by filtering the sample of individuals that were recorded as sexually mature (pregnant, ovulating, lactating, etc). These samples were prioritized as being reported as mature is an indicator that biologists or trained personnel carefully examined the catch, resulting in more accurate measurements. Since this cannot be replicated for the male catches, the male length frequencies were dissected in normal probability plots to obtain the component curves for the mature population from inflection points (Harding, 1949).

For accurate results, the mean length of sexual maturity (*L_sm_*) was used as a starting value that was adjusted slightly for fitting the component curves. While early studies for *P. macrocephalus* interpreted sexual maturity in males to occur at 9.5-12.5 m (Aguayo, 1963; Clarke, 1956; Matthews, 1938; Nishiwaki, 1955; Nishiwaki et al., 1958), the estimated *L_sm_*at 13.7-14.4 m is likely more accurate and was selected for this analysis. This size coincides with the histological maturation at the periphery of the testes, sudden increase in testes mass/volume, deceleration of body growth prior to physical maturity, elevation of breeding status, and the transition to solitude (Best, 1969, 1970, 1979; Clarke & Paliza, 1988; Gambell, 1972; Kato, 1984).

The y-intercept and slope of these linear segments in the probability plots were used to estimate the *L*_mean_ and SD, respectively. In cases where too few females were recorded as mature (<30), the total female catch was graphically analyzed in the same manner as males. The mean lengths for each sex between regions were compared using the two-sample *t*-test (Zar, 2010).

As a simple reference for identifying exceptional lengths, the expected upper bound for the maximum value within a normally-distributed sample of *n* (Equation 4) is compared against potentially unreliable values within the whaling data (Kamath, 2015). For *P. macrocephalus*, *n* is the total number of whales in the entire IWC catch database that were greater than or equal to the *L*_sm_ and rounded to the nearest thousand (*Y* = deviation from mean length).

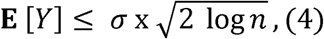

Since natural variation could still allow individuals to exceed the expected maximum length from Equation 4, reliable estimates for the *L*_max_ of *P. macrocephalus* will be determined using a combination of the verified measurements in literature and the largest sizes recorded across different expeditions in the commercial data. This helps to account the possible influence of biases in measuring technique.

Because the IWC catch database is not comprehensive for historical catches of delphinids and ziphiids, additional data from literature (Benjaminsen, 1972; Christensen, 1975, 1984; Degerbøl, 1940; Fernandez et al., 2014; Grove et al., 2020; Omura et al., 1955; Reeves et al., 1993; Thompson, 1846) and museum records (Natural History Museum, 2018; Orrell & Informatics Office, 2024) were sourced to calculate the *L*_mean_ for *O. orca* and the two ziphiid species. For the two ziphiid species, graphical analysis was employed to calculate the *L*_mean_ in both sexes.

Due to both the mixture of immature individuals and multiple ecotypes within historical male catches, only the *L*_mean_ for females of the North Atlantic (NA) Type 1 ecotype in Norwegian catches (*n* = 51) and Antarctic (ANT) Type A ecotype (*n* = 125) were precisely calculated. While ovulations begin at sexual maturity, females typically bear viable calves around the age of physical maturity (Fearnbach et al., 2011; Kotik et al., 2023; Olesiuk et al., 1990). Therefore, the mean lengths reported for adult females accompanied by calves are treated as estimates of *L*_∞_.

## 3 Review of size distributions and body mass relationships

The *L*_mean_, *L*_max_, and corresponding masses of four odontocete species are summarized in Table 1.

**Table 1:**
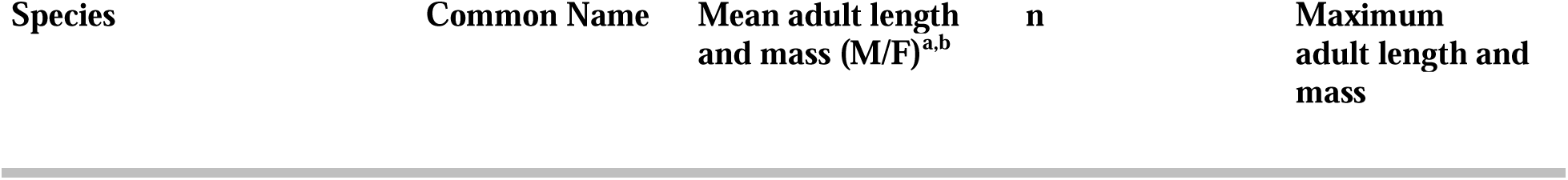
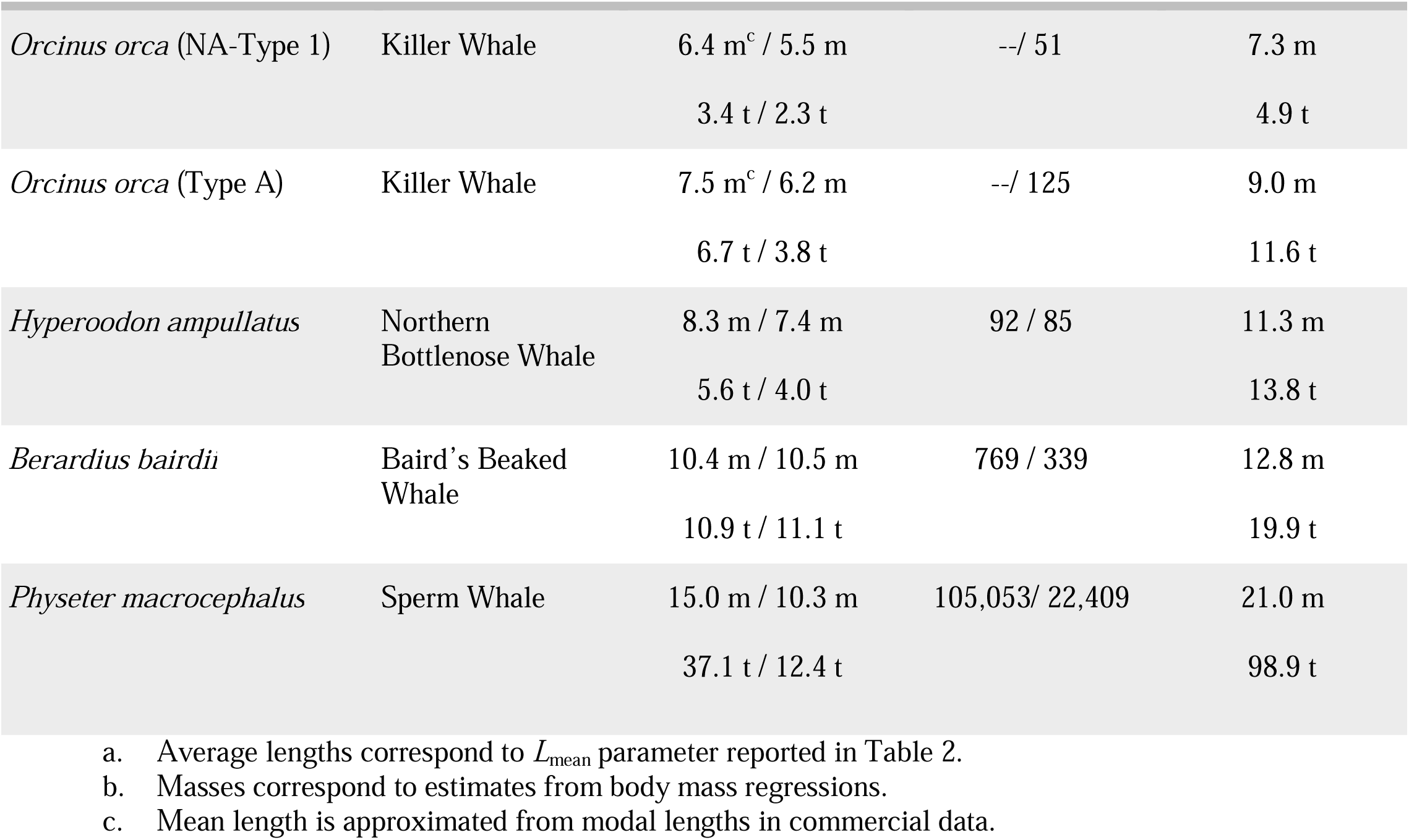
Summary of length and mass for large odontocetes.

| Species | Common Name | Mean adult length and mass (M/F)<sup>a,b</sup> | n | Maximum adult length and mass |
| --- | --- | --- | --- | --- |
| *Orcinus orca* (NA-Type 1) | Killer Whale | 6.4 m<sup>c</sup> / 5.5 m<br>3.4 t / 2.3 t | --/ 51 | 7.3 m<br>4.9 t |
| *Orcinus orca* (Type A) | Killer Whale | 7.5 m<sup>c</sup> / 6.2 m<br>6.7 t / 3.8 t | --/ 125 | 9.0 m<br>11.6 t |
| *Hyperoodon ampullatus* | Northern Bottlenose Whale | 8.3 m / 7.4 m<br>5.6 t / 4.0 t | 92 / 85 | 11.3 m<br>13.8 t |
| *Berardius bairdii* | Baird's Beaked Whale | 10.4 m / 10.5 m<br>10.9 t / 11.1 t | 769 / 339 | 12.8 m<br>19.9 t |
| *Physeter macrocephalus* | Sperm Whale | 15.0 m / 10.3 m<br>37.1 t / 12.4 t | 105,053/ 22,409 | 21.0 m<br>98.9 t |

### Orcinus orca

Modern photogrammetry studies and the re-analysis of historical data have enabled experts to observe significant differences in adult size across multiple sympatric ecotypes of *O. orca* (Best et al., 2010; Durban et al., 2021; Fearnbach et al., 2011; Foote et al., 2009; Groskreutz et al., 2019; Kotik et al., 2023; Pitman et al., 2007). The two ecotypes listed in Tables 1 & 2 reflect the consistent difference in average and maximum adult size observed between mammal-specialists (Eastern North Pacific (ENP) Transient, NA Type 2, ANT Type A, ANT Type B1) and piscivorous/generalist ecotypes (ENP Resident, ENP Offshore, NA Type 1, ANT Type B2, and ANT Type C). A polymodal distribution for mature Antarctic females within the IWC dataset (Figure S1) appears to be a mixture of the Type A ecotype and the much smaller Type C as suspected in previous examinations (Mikhalev et al., 1981; Pitman & Ensor, 2003).

Published body mass regressions for *O. orca* from ENP live-capture operations (Bigg & Wolman, 1975), Soviet whaling (Mikhalev, 2019), and biological studies on the Type A ecotype in southern Africa (Best, 2008) are provided in Table 3. Figure 1A and Figure S2 compare the 51 whole weights collected from live-capture records and necropsies for the resident (*n* = 28), transient (*n* = 13), and NA Type 1 ecotypes (*n* = 10) after the removal of two outliers (Hoyt, 1990; Yamada et al., 2007). The AIC selection for body mass regressions strongly favored a model separating the ecotypes (AICc = -109.91, weight = 0.95) rather than a pooled sample (ΔAICc = 6.05, weight = 0.05). The curves for the NA Type 1 and resident ecotypes were nearly identical while transients were notably heavier, as in the case for the South African and the Soviet curves.

**Figure 1:**
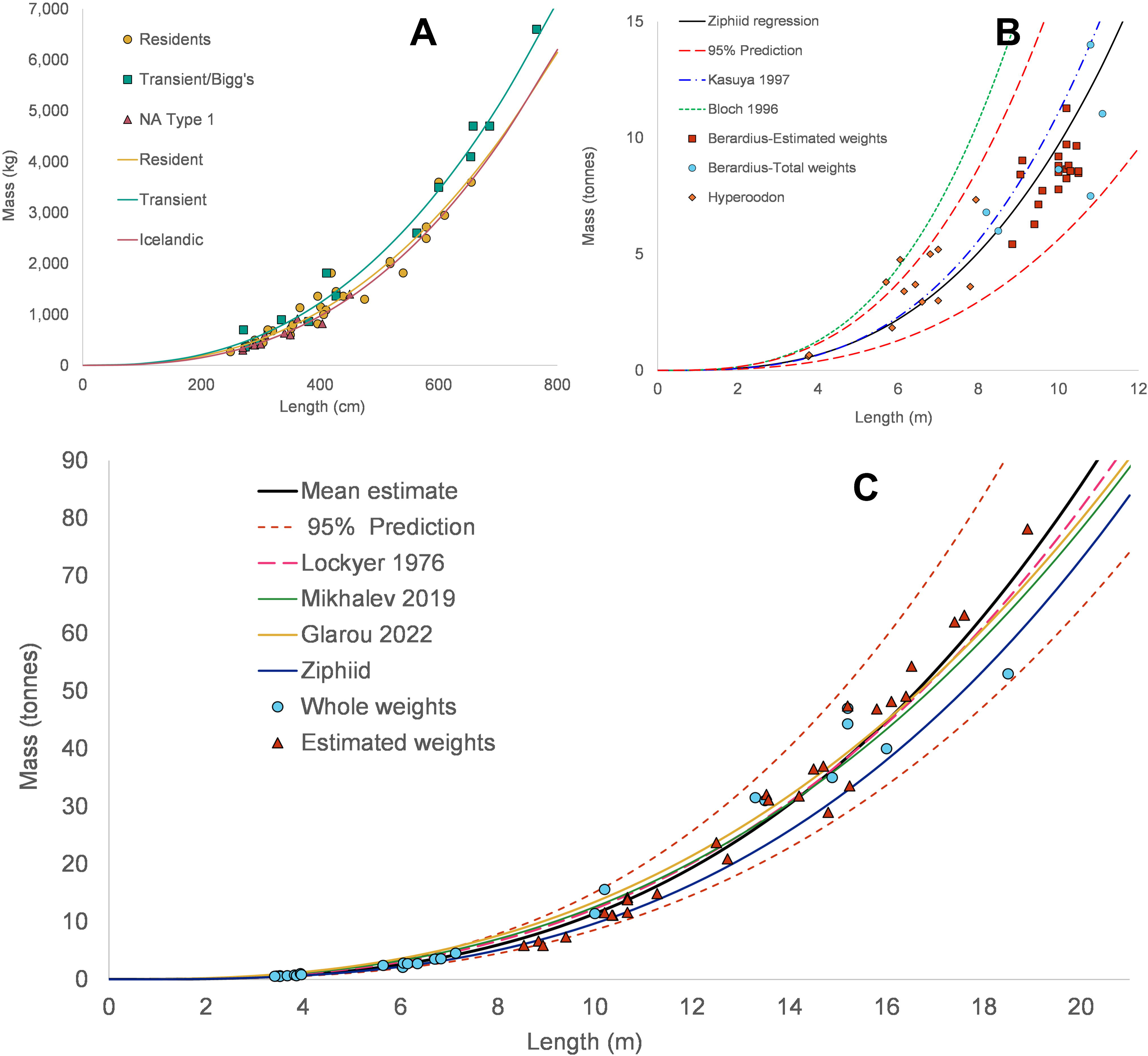
Length-weight relationships in large odontocetes. Legend: Length-weight curves for *O. orca* (A), ziphiids (B), and *P. macrocephalus* (C).

A 9.5 m male (presumably a transient) reported within Japanese fishery data is the largest specimen noted in literature (Nishiwaki & Handa, 1958). The heaviest mass ever published for *O. orca* was a record of 9.95 tonnes for an 8.6 m Antarctic male (Sleptsov, 1965). The intact weight would have been 10.5 tonnes with at least a 5% correction for the body fluids (Ridgway & Johnston, 1966). The live-capture and Soviet whaling regressions predict a 6.5 m fish-specialist weighing 3.6 tonnes, a 7.5 m mammal-specialist weighing 6.7 tonnes, and a 9.5 m mammal-specialist weighing 13.6 tonnes.

### Hyperoodon ampullatus *&* Berardius bairdii

*H. ampullatus* and *B. bairdii* represent both the largest and most well-studied ziphiid species. *H. ampullatus* exhibits a male-biased sexual dimorphism (Table 2 & Figure S3), with maximum male and female lengths of 11.3 m and 8.7 m, respectively (Degerbøl, 1940; Thompson, 1846). *B. bairdii* displays the opposite pattern (Table 2 & Figure S4), with males and females respectively reaching 11.9 m and 12.8 m (Nishiwaki & Oguro, 1971).

**Table 2:**
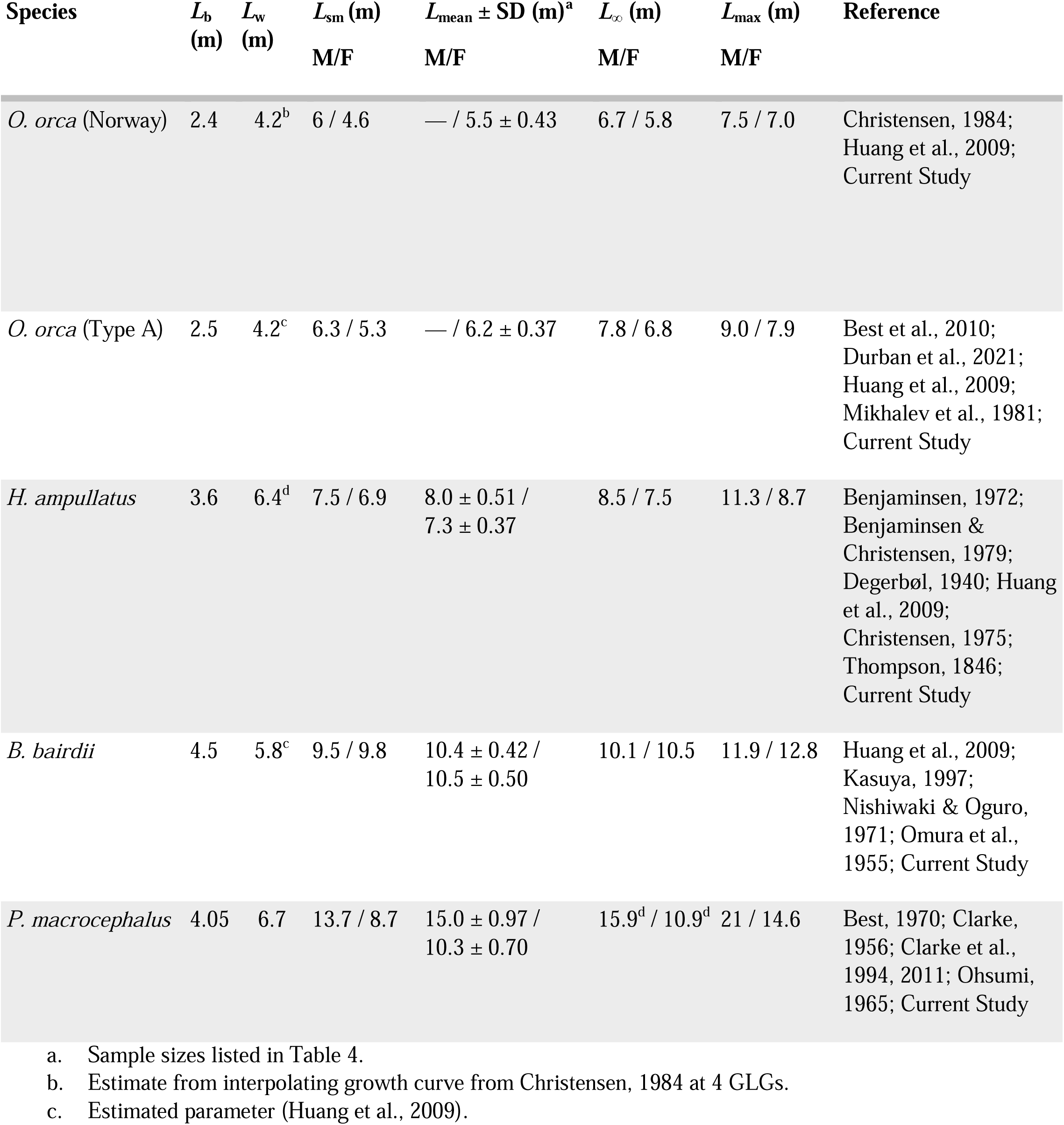
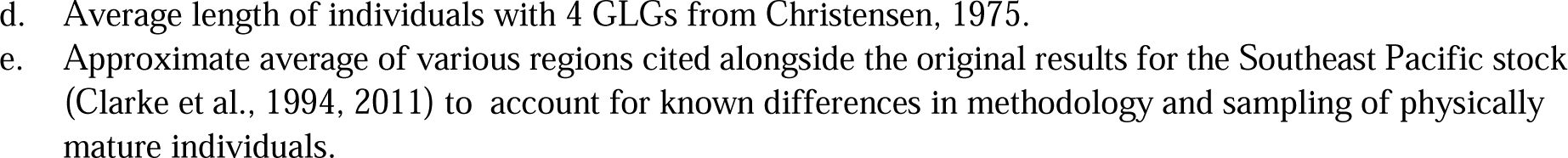
Size parameters for large odontocetes.

Total body masses for 20 specimens have been published between *Hyperoodon* (*n* = 14) and *Berardius* (*n* = 6). The heaviest masses recorded for *H. ampullatus* and *B. bairdii* were a 7.34-tonne male and a 14-tonne lactating female, respectively (Fernandez et al., 2014; Kasuya, 1997). The published body mass models for both species (Table 3) tend to overestimate most of the published data (Figure 1B). These formulas were probably skewed by either seasonal fattening or imprecise measuring of product yields during the Faroese harvests (Bloch et al., 1996) and limited sampling (Kasuya, 1997).

**Table 3:**
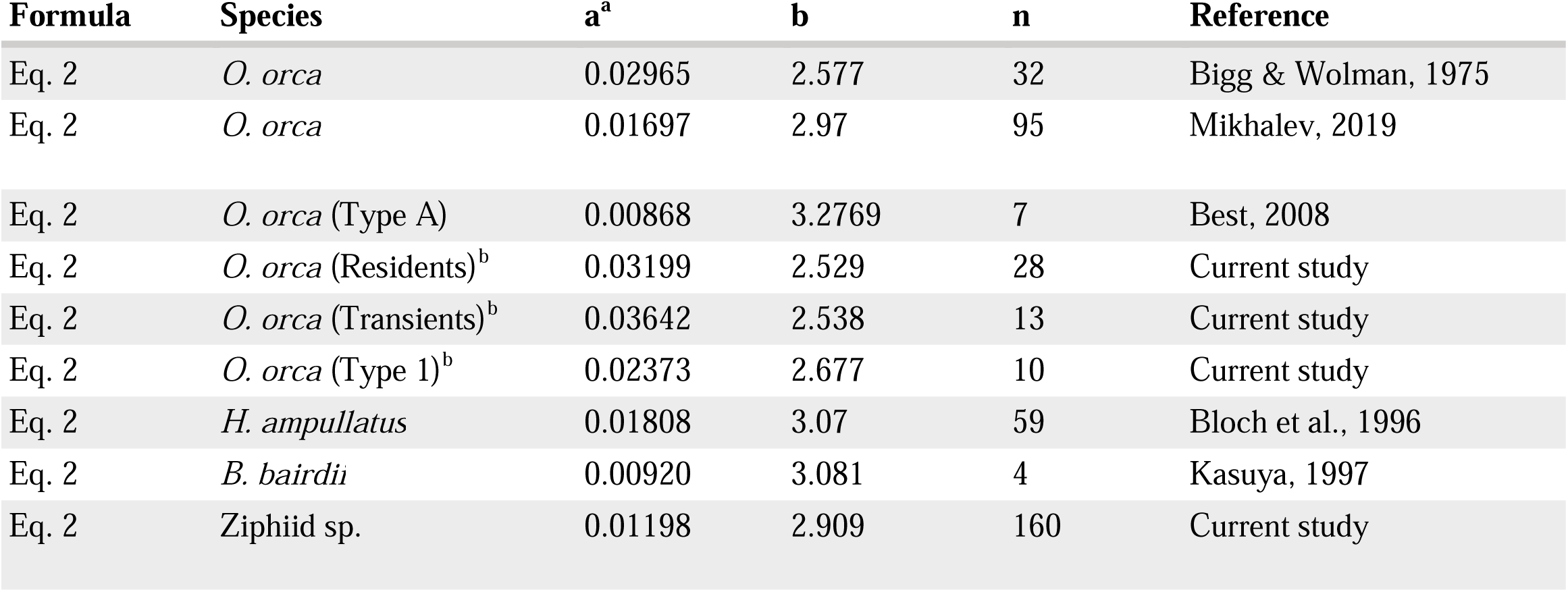
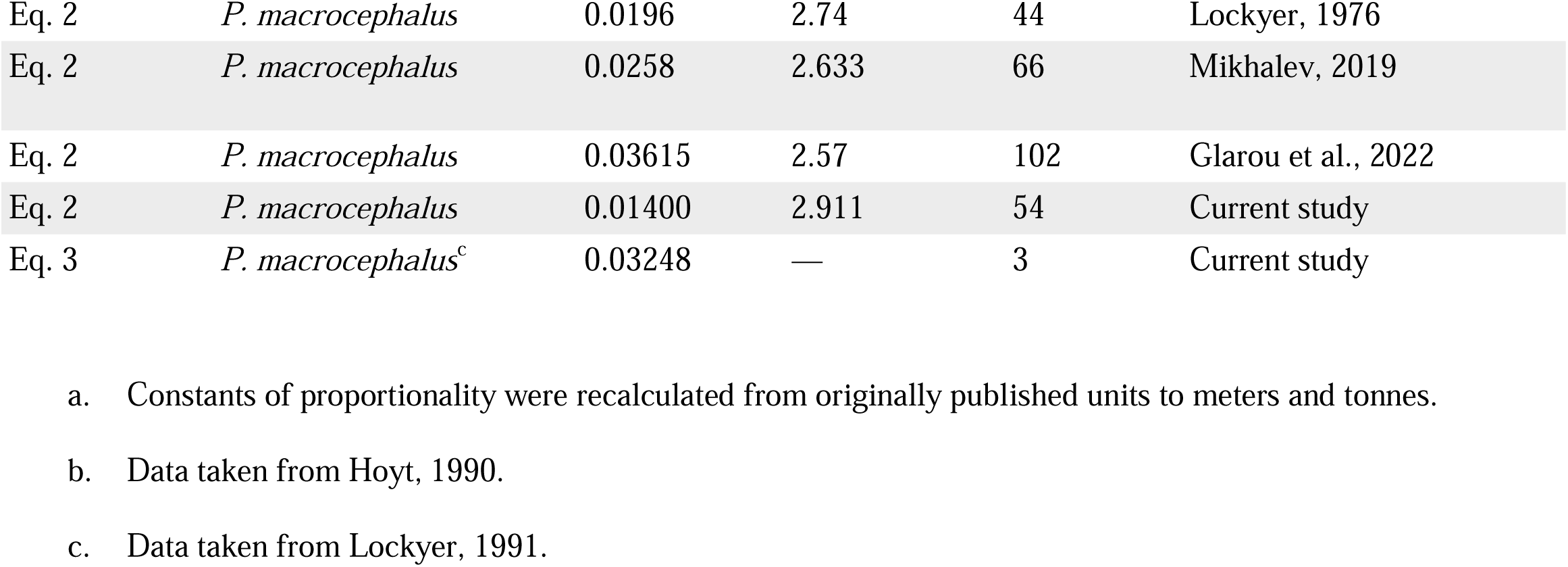
Body mass regression parameters for large odontocetes.

| Formula | Species | a<sup>a</sup> | b | n | Reference |
| --- | --- | --- | --- | --- | --- |
| Eq. 2 | *O. orca* | 0.02965 | 2.577 | 32 | Bigg & Wolman, 1975 |
| Eq. 2 | *O. orca* | 0.01697 | 2.97 | 95 | Mikhalev, 2019 |
| Eq. 2 | *O. orca* (Type A) | 0.00868 | 3.2769 | 7 | Best, 2008 |
| Eq. 2 | *O. orca* (Residents)<sup>b</sup> | 0.03199 | 2.529 | 28 | Current study |
| Eq. 2 | *O. orca* (Transients)<sup>b</sup> | 0.03642 | 2.538 | 13 | Current study |
| Eq. 2 | *O. orca* (Type 1)<sup>b</sup> | 0.02373 | 2.677 | 10 | Current study |
| Eq. 2 | *H. ampullatus* | 0.01808 | 3.07 | 59 | Bloch et al., 1996 |
| Eq. 2 | *B. bairdii* | 0.00920 | 3.081 | 4 | Kasuya, 1997 |
| Eq. 2 | Ziphiid sp. | 0.01198 | 2.909 | 160 | Current study |
| Eq. 2 | *P. macrocephalus* | 0.0196 | 2.74 | 44 | Lockyer, 1976 |
| Eq. 2 | *P. macrocephalus* | 0.0258 | 2.633 | 66 | Mikhalev, 2019 |
| Eq. 2 | *P. macrocephalus* | 0.03615 | 2.57 | 102 | Glarou et al., 2022 |
| Eq. 2 | *P. macrocephalus* | 0.01400 | 2.911 | 54 | Current study |
| Eq. 3 | *P. macrocephalus*<sup>c</sup> | 0.03248 | — | 3 | Current study |

a. Constants of proportionality were recalculated from originally published units to meters and tonnes.
b. Data taken from Hoyt, 1990.
c. Data taken from Lockyer, 1991.

For this review, a composite regression for Ziphiidae was calculated using the data collected for *Hyperoodon*, *Berardius*, and 121 observations from 15 other ziphiid species to improve length coverage (Table 3, Figure 1B, & Figure S5). Within this sample includes estimated total body masses from the muscle yields for 19 *B. bairdii* specimens ranging from 8.9-10.5 m (Balcomb, 1989). These yields were divided by 0.38, the proportional weight of the muscle from a piecemeal weighing of a 7.5-tonne non-pregnant specimen (Tomilin, 1967). While it was reasonably assured that nearly all other data compiled were direct measurements, it was not always discernible whether some included corrections for body fluids.

Despite the inclusion of 18 ziphiid species in total, the correlation for the linear regression was fair (*n* = 160, r^2^ = 0.955), likely owing to similarity in overall body shape among ziphiids. While AIC model selection for the entire sample strongly preferred separating the species (AICc =-239.1, weight = 1), a pooled model was slightly preferred when data for *B. bairdii* was omitted (AICc = -185.11, weight = 0.55). This result is easily attributable to the poor length coverage within the *B. bairdii* data, as the slope (b = 1.85) was unrealistic given the approximately cubic relationship between TBL and weight in cetaceans. Limited data on TBL and weight of individual ziphiids will continue to limit inter- and intraspecific morphological studies of growth.

### Physeter macrocephalus

Despite whale-stretching causing an artificial peak at 10.7 m, the length distributions for mature females captured between 1937-1971 resembled those captured after 1972 and in illegal catches by the Soviet Union that ignored length restrictions. Graphical analysis of the pooled global male dataset (*n* = 232,975) estimated that 45% (*n* =105,053) of all males were sexually mature (Figure S6). This differs slightly from the sum of mature males in Table 4 (*n* =109,739), as the exact inflection point varied slightly between regions. While all regional mean length differences ≥ 0.1 m were statistically significant (*t-*test, *p* < 0.05), the intraregional shifts in means between pre- and post-IOS periods overlapped with most of the interregional variation.

**Table 4:**
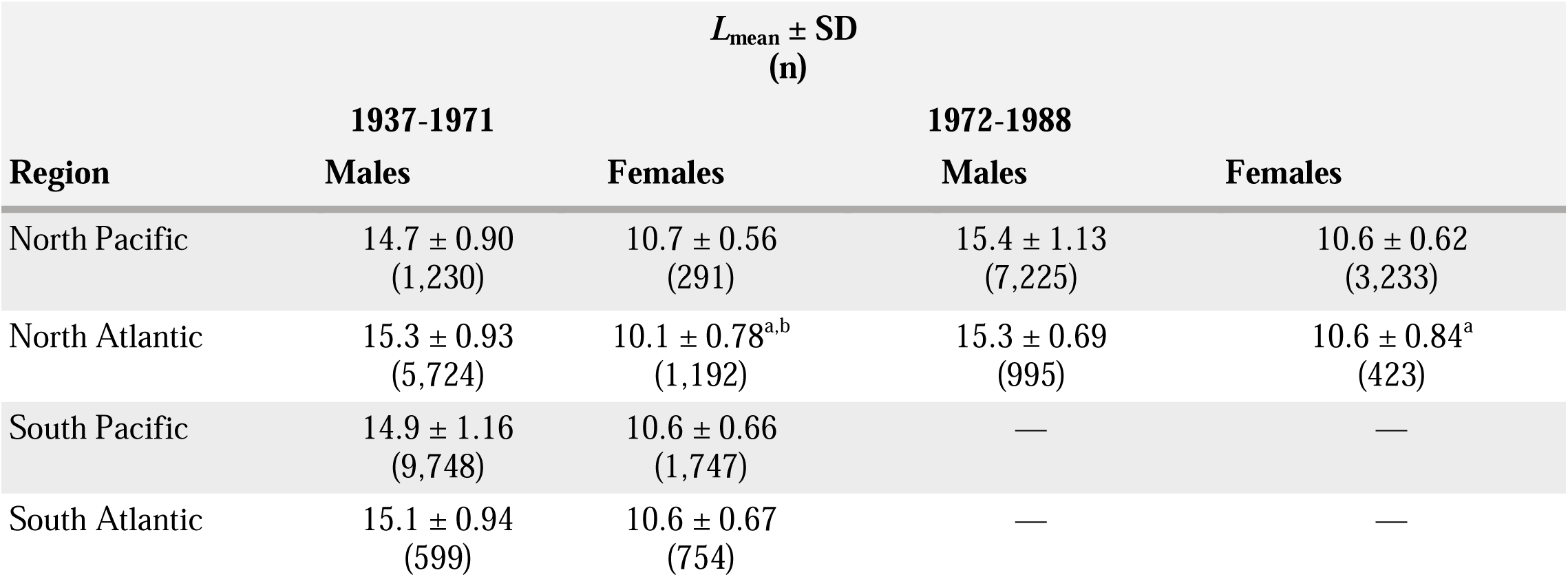
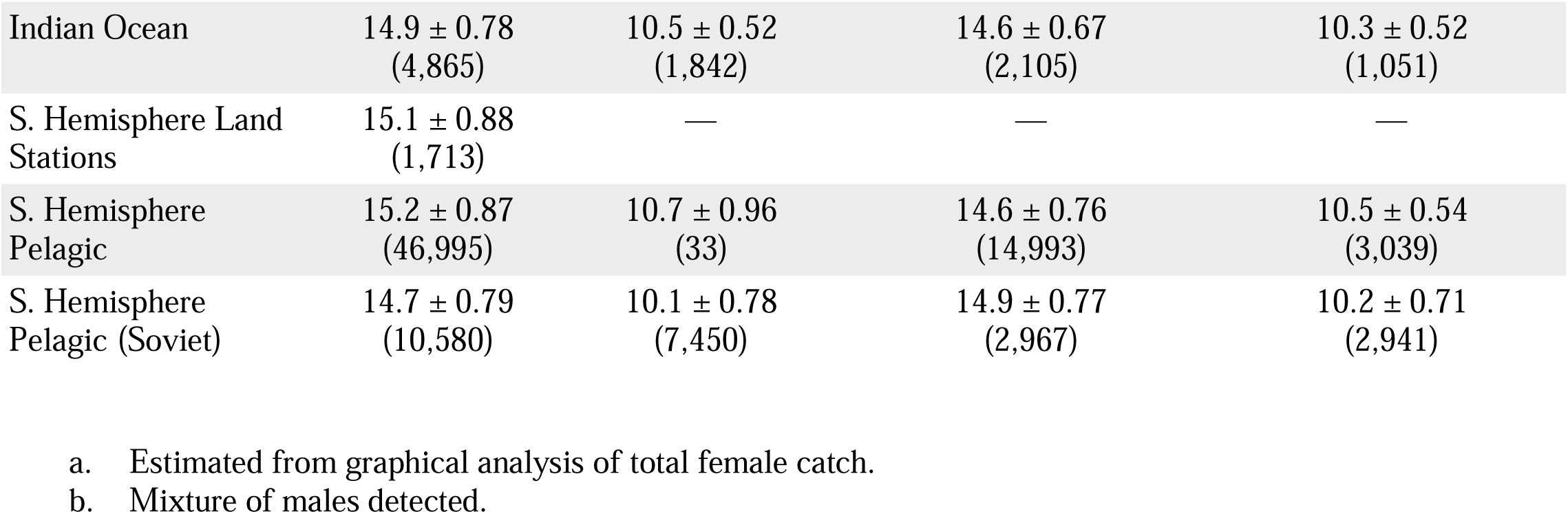
Regional comparison of mean adult length (*L*_mean_) of *P. macrocephalus*.

| Region | $L_{\text{mean}} \pm \text{SD}$<br>(n) | | | |
| --- | --- | --- | --- | --- |
|  | 1937-1971<br>Males | Females | 1972-1988<br>Males | Females |
| North Pacific | 14.7 ± 0.90<br>(1,230) | 10.7 ± 0.56<br>(291) | 15.4 ± 1.13<br>(7,225) | 10.6 ± 0.62<br>(3,233) |
| North Atlantic | 15.3 ± 0.93<br>(5,724) | 10.1 ± 0.78<sup>a,b</sup><br>(1,192) | 15.3 ± 0.69<br>(995) | 10.6 ± 0.84<sup>a</sup><br>(423) |
| South Pacific | 14.9 ± 1.16<br>(9,748) | 10.6 ± 0.66<br>(1,747) | — | — |
| South Atlantic | 15.1 ± 0.94<br>(599) | 10.6 ± 0.67<br>(754) | — | — |
| Indian Ocean | 14.9 ± 0.78<br>(4,865) | 10.5 ± 0.52<br>(1,842) | 14.6 ± 0.67<br>(2,105) | 10.3 ± 0.52<br>(1,051) |
| S. Hemisphere Land Stations | 15.1 ± 0.88<br>(1,713) | — | — | — |
| S. Hemisphere Pelagic | 15.2 ± 0.87<br>(46,995) | 10.7 ± 0.96<br>(33) | 14.6 ± 0.76<br>(14,993) | 10.5 ± 0.54<br>(3,039) |
| S. Hemisphere Pelagic (Soviet) | 14.7 ± 0.79<br>(10,580) | 10.1 ± 0.78<br>(7,450) | 14.9 ± 0.77<br>(2,967) | 10.2 ± 0.71<br>(2,941) |

- a. Estimated from graphical analysis of total female catch. - b. Mixture of males detected.

Only a few females exceeding 12 m have ever been verified in literature (Berzin, 1964; Clarke, 1956), the largest was a 14.3 m specimen examined near the Kuril Islands in 1948 (Sleptsov, 1955). Records male-like size distributions for females from the unfiltered total catches (McClain et al., 2015) are likely artefacts of whalers misidentifying the sex of males with retracted penises (Berzin, 1964; Matthews, 1938). A small percentage of misreported males were also apparent within the total catch of North Atlantic females in the pre-IOS dataset (Figure S6). As expected, sex misidentification was mitigated for the commercial dataset of females recorded as sexually mature. The largest in the current dataset were two pregnant females measuring 14.6 m were reported by Spanish land stations in 1950.

While most literature cites 18.3 m as the maximum length for males (Rice, 1989), skeletal examination and aging methods suggested that multiple bulls measuring 18-18.6 m were physically immature (Andrews, 1916; Boschma, 1938; Clarke, 1956; Kasuya, 1991; Mitchell & Kozicki, 1984). A close review of existing stranding records and mounted skeletons (Figure S7&S8) verified bulls measuring 18.9-19.2 m (Newman, 1910; Wood, 1972). The largest bull to be measured and photographed was reportedly 20.7 m and captured in 1903 off Ronas Voe (Haldane, 1904; Millais, 1906). Measurements taken from the photograph of the anterior body appear to substantiate the recorded length (Figure S8 & Table S1). In 1950, a 20.8 m bull was also recorded off the Kuril Islands, which is among the reliable portions of the pre-IOS Soviet data (Ivashchenko et al., 2014) that were excluded in this analysis. This same whale is canonically cited in other literature (Wood, 1972; Zenkovich, 1954).

The largest individual within the IWC dataset for the current analysis was a 21.0 m bull captured off Peru by a British whaling flotilla in 1937. The next largest were 13 bulls of similar size (20.1-20.7 m) taken by Chilean land stations in the Southeast Pacific and U.S. land stations in the North Pacific between 1938-1955. It appears that 21 m is the largest reliable size within the commercial data.

A 24 m bull captured by Chilean fleets in 1933 was previously reported as the record size for *P. macrocephalus* (McClain et al., 2015). Morphometric analyses (Figure S7) on other bulls allegedly measuring 24-27.4 m (Starbuck, 1878; Wood, 1982) suggest that such TBLs were not standard measurements. Accounting for excluded catches (falsified and pre-1937) along with the 100,000-110,000 mature bulls reported in Tables 1 & 2 would suggest around 150,000 mature bulls were captured throughout the 20^th^ century. At this population size, the maximum size reaching 9 meters (9.3 SDs) beyond the global mean length (Table 2) would seem unrealistic.

A maximum length of 21 m for bulls is more consistent with the deviations for the largest reliably measured females within literature and commercial datasets (∼ 6 SD) and is much closer to, but still exceeding, the upper bound approximation (Equation 4) of 19.7 m when n=150,000. While this discrepancy could be owed to errors from the measuring technique or the condition of the carcass (Clark, 1983), the effect of nourishment on the adult size and growth of sperm whales (Clarke et al., 1994; Kahn et al., 1993; Kasuya, 1991) could likely facilitate the occurrence of outliers.

The heaviest whole weight recorded for *P. macrocephalus* was 53 tonnes for an 18.5 m bull (Boschma, 1938; Deinse, 1946). An 18.1 m bull with a 57.1-tonne product yield would have weighed at least 64.9 tonnes with a 12% correction for blood loss applied (Tomilin, 1967). Intact weights from 29 whales were compiled for this review. Morphometric data for three of these whales were used to develop the RW model (*n* = 3, r^2^ = 0.998). The RW model for *P. macrocephalus* was tested against four bulls with fluid-corrected weights of 25.8-60.6 tonnes (Zenkovich, 1952), whose maximum girths were converted from the corresponding body widths (Glarou et al., 2022). The correlation for the predicted masses of these four bulls (r^2^= 0.985) compared favorably to those other species (George, 2009; Sumich et al., 2013). The accuracy of the RW model over such an extensive size range warranted its use to estimate the body masses of 29 additional whales from length and girth measurements.

The RW model predicted a weight of 100.1 tonnes for the Ronas Voe bull (Haldane, 1904), although the largest estimate from verified measurements was 78.1 tonnes for an 18.9 m bull from Port Arthur, Texas (Newman, 1910). Within the overlapping length range (≥ 10 m), AIC model selection preferred pooling both estimated and direct weights for one regression (AIC= - 86.81, weight= 0.77). The resulting model, which excludes the Ronas Voe bull, is presented in Table 2 and Figure 1C along with other published weight curves. Piecemeal weight curves (Lockyer, 1976; Mikhalev, 2019) have been adjusted upward by 12%.

The linear regression for the pooled sampled had a strong correlation (Figure S9, n=54, r^2^ = 0.992) and closely agreed with the curves derived from piecemeal weights and photogrammetry (Glarou et al., 2022; Lockyer, 1976; Mikhalev, 2019). While AIC selection preferred separate weight relationships for ziphiids and *P. macrocephalus* (AICc= -346.52, weight= 0.99), comparing the clades in a multiple regression analysis did not detect a significance difference for either the slopes (*p* = 0.975) or the intercepts (*p* = 0.340). The whole weight curve predicts *P. macrocephalus* weighing 15.1 tonnes (95% PI: 11.3-20.0 tonnes) at 11 m, 37.1 tonnes (95% PI: 27.9-49.3 tonnes) at 15 m, and 98.9 tonnes (95% PI: 74.1-131.9 tonnes) at 21 m.

The record oil yield reported for a single sperm whale in the 19^th^ century was 168 barrels/17.5 tonnes (Banks, 1911), which typically represents 18-24% (mean = 20.2%) of a whale’s piecemeal weight based on the known lipid content of blubber and spermaceti (Berzin, 1972; Heptner et al., 1988; Ohno & Fujino, 1952; Omura, 1950; Zenkovich, 1937, 1954). The respective percentages predict an intact weight of 83.4-111.1 tonnes (mean=99.2 tonnes), converging with both the whole weight curve’s estimate for a 21 m bull and the RW model’s prediction for the Ronas Voe bull. There appears to be little evidence that *P. macrocephalus* attained a greater maximum size during the 19^th^ century.

### Relationships between length parameters

It is already recognized that marine mammals sexually mature at around 85% the TBL of physical maturity (Laws, 1956), which is obeyed by data reported in Table 2 (mean=85.4%). Across the four species reviewed (Table 5), *L*_sm_ equaled 89.8% of the *L*_mean_, and the *L*_mean_ equaled 96.2% of the *L*_∞_. For the more highly sampled species, *B. bairdii* and *P. macrocephalus*, the CV was very similar between sexes. The CV is notably higher in *P. macrocephalus*, which also exhibits a greater relative difference between *L*_mean_ and *L*_∞_.

**Table 5:** Intraspecific relationships between *L*_sm_, *L*_mean_ , *L*_∞_, and CV in large odontocetes.

| Species | $L_{\text{sm}}/L_{\text{mean}}$ (%)<br>M/F | $L_{\text{mean}}/L_{\infty}$ (%)<br>M/F | CV (%)<br>M/F | n<br>M/F |
| --- | --- | --- | --- | --- |
| *O. orca* |  |  |  |  |
| Type 1 | — / 83.6 | — / 94.8 | — / 7.82 | — / 51 |
| Type A | — / 85.5 | — / 91.2 | — / 5.79 | — / 125<sup>a</sup> |
| *H. ampullatus* | 93.4 / 94.5 | 94.1 / 97.3 | 6.38 / 5.07 | 92<sup>a</sup> / 85<sup>a</sup> |
| *B. bairdii* | 92.2 / 93.3 | 102<sup>b</sup> / 100 | 4.04 / 4.76 | 769<sup>a</sup> / 339<sup>a</sup> |
| *P. macrocephalus* | 91.3 / 84.5 | 94.3 / 94.5 | 6.47 / 6.80 | 105,053<sup>a</sup> / 22,409 |

a. Approximate sample size within component curve.
b. This result is likely due to the difference in the level of measuring precision from commercial data. Should be treated as 100% like in females.

## 4 Discussion

Some species or stocks of large odontocetes, such as ziphiids or fossil taxa (Lambert et al., 2010; MacLeod, 2006), are known from only a scant number of measurements for which preliminary values for the *L*_sm_ or *L*_∞_ are available. Therefore, clarifying a consistent relationship between these parameters with the *L*_mean_ and coefficient of variation (CV) in related species of similar size would provide assumptions for modeling entire adult size distributions from only an estimate for the *L*_sm_ or *L*_∞_.

The differences in the adult sizes parameters observed between *P. macrocephalus* and *B. bairdii* were consistent with previous research suggesting that growth being more prolonged in *P. macrocephalus* (Clarke et al., 1994, 2011; Kasuya, 1997). There might be some uncertainty in comparing the *L*_mean_ and *L*_∞_ for *B. bairdii* due to differences in the precision of measurements between commercial data and that of scientists (Kasuya, 1997). Nonetheless, this would similarly apply to *P. macrocephalus* (Kasuya, 1991). Limited sample sizes prevent confident interpretations regarding the CV for *O. orca* and *H. ampullatus*. The limitations present in this current analysis, other than acquiring more data for the current species, can be addressed by extending analyses towards the abundant commercial data available for mysticetes.

While regional differences in *L*_mean_ were statistically significant for *P. macrocephalus*, they did not exceed the expected intraregional variation. Since *P. macrocephalus* may continue to physically grow for decades after becoming sexually mature, the *L*_mean_ is likely sensitive to differences in the natural age composition or catch selectivity. Therefore, caution should be placed in assuming a diagnostic morphological difference between populations without data on age or physical development. As it currently stands, asymptotic size estimates appear to be largely congruent between regions when accounting for known differences in estimation methods and the limited sampling of physically mature individuals for certain regions (Clarke et al., 1994, 2011).

Differences in weight relationships between the *O. orca* ecotypes reviewed here were not detected in a previous analysis (Williams et al., 2011), which is likely due to their dataset lacking adult-sized transients. While ecotypes were not specified for the Soviet weight curves, marine mammals were observed in the stomachs in the majority of the catch (Shevchenko, 1975; Sleptsov, 1965). Mammal-eating killer whales being proportionally heavier is consistent with photogrammetric measurements of body width in the ENP and Antarctic (Durban et al., 2021; Kotik et al., 2023). The weight data for residents was collected 20 years prior to the declines in Chinook salmon abundance (Ford et al., 2010), suggesting that mammal-eating killer whales are naturally more robust even when compared to resident populations that were in healthier condition. This may be an important factor to consider in future morphometric studies of killer whales.

Pooling all the data for Ziphiidae returned a weight relationship like that of *P. macrocephalus*. AIC model selection detected a difference in the linear models for the two clades, though this may be attributable to an undetermined number of uncorrected piecemeal weighings within the ziphiid sample. Multiple regression analysis was at least able to support that the slopes were nearly identical. This morphological similarity may reflect their convergence towards a deep-diving, teuthophagous ecology (Kenny et al., 1985; Rice, 1989).

## 5 Conclusions

This review on the size of large cetaceans has compiled a large breadth of morphometric data for postnatal growth, adult variation, and length-weight relationships. Existing literature was heavily covered to summarize familiar data, bring old records out from obscurity, and provide relationships between adult size parameters. This review may aid advancing morphometric analyses that emphasize the importance of intraspecific variation.

### Data availability

Supplementary figures, code, and morphometric data acquired from literature and museum records can be acquired upon contact with the corresponding author. All whaling data is available from the International Whaling Commission upon request to this address

IWC contact page: https://iwc.int/contact

## Acknowledgements

The author expresses appreciation towards Dr. Sally Mizroch, Dr. Trevor Branch, Dr. Fredrik Christiansen, Dr. Yulia Ivashchenko, Dr. Phil Clapham, Dr. James Mead, Seán O’Callaghan, and Samuel Dunford for both their correspondence and assistance in acquiring important sources for this paper. The author thanks the Bayworld Museum’s marine mammal curator, Dr. Greg Hofmeyr, and collection manager, Dr. Sibu Ngquana, for providing jaw length data for *P. macrocephalus*. The author also acknowledges Ayla Parker for creating the 3D sperm whale model used in the photogrammetric analysis in the supplementary material.

