## Supplementary File 1 for "A summary of intraspecific size variation for large odontocetes"

### **A summary of intraspecific body size variation of large odontocetes. Supplementary File 1**

Author: Joseph McClure*

*Correspondence

564 East McIver Road, Florence, South Carolina 29506

#### Figure S1. Length distribution and normal probability plot of Soviet catch of mature female killer whales in the Southern Hemisphere. 73% of the catch contains the larger ecotype (likely Type A) and 27% correspond to Type C.

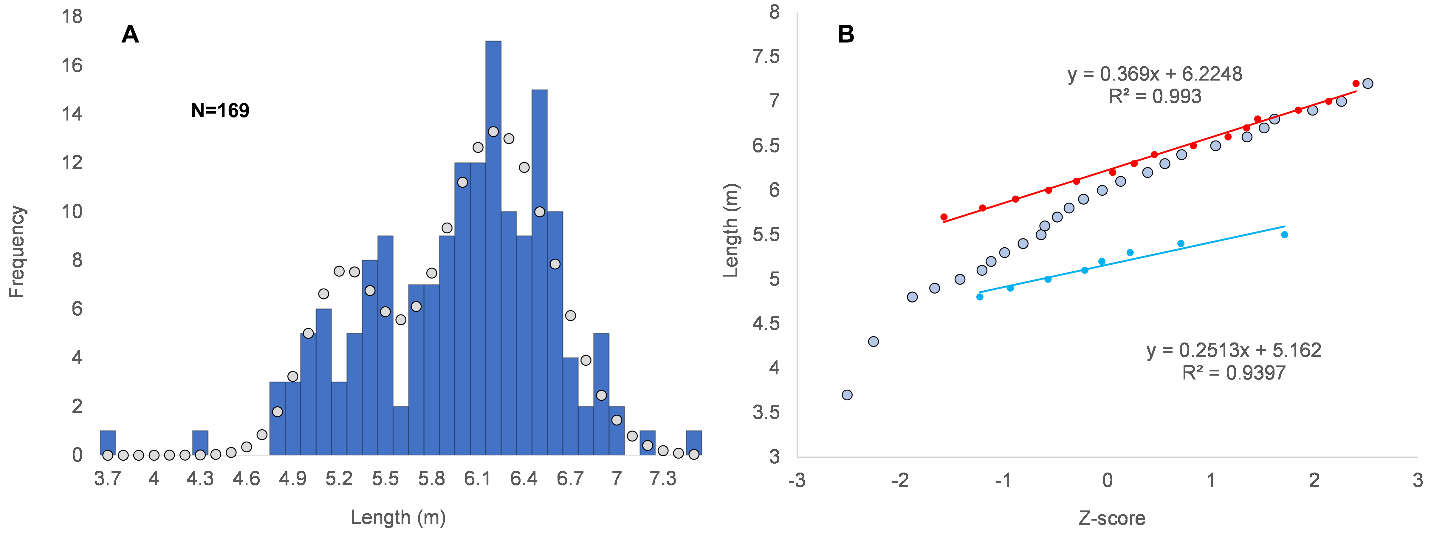

#### **Figure S2.** Linear regressions for log-transformed length and masses for resident (y=2.529x -3.5502, n=28, r^2^=0.938), transient (y=2.538x - 3.516, n=13, r^2^=0.950), and NA Type 1 Killer whales (y=2.677x - 3.978, n=10, r^2^=0.916). Two Icelandic outliers (270cm/760kg; 394cm / 600kg) were omitted. See Supplementary File 2 for data and File 3 for references.

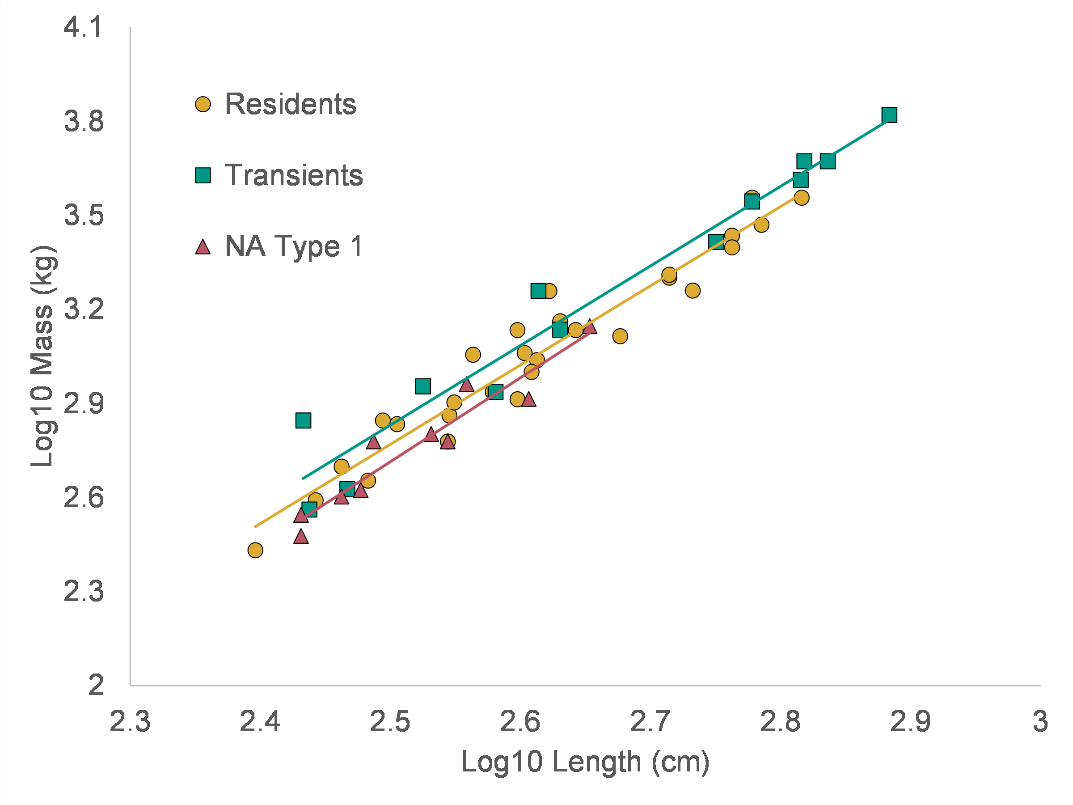

#### **Figure S3.** Length distributions and normal probability plots for female (A&B) and female (C&D) northern bottlenose whales. Approximately 45% of males and 52% of females were mature. See supplementary File 2 for data and File 3 for references.

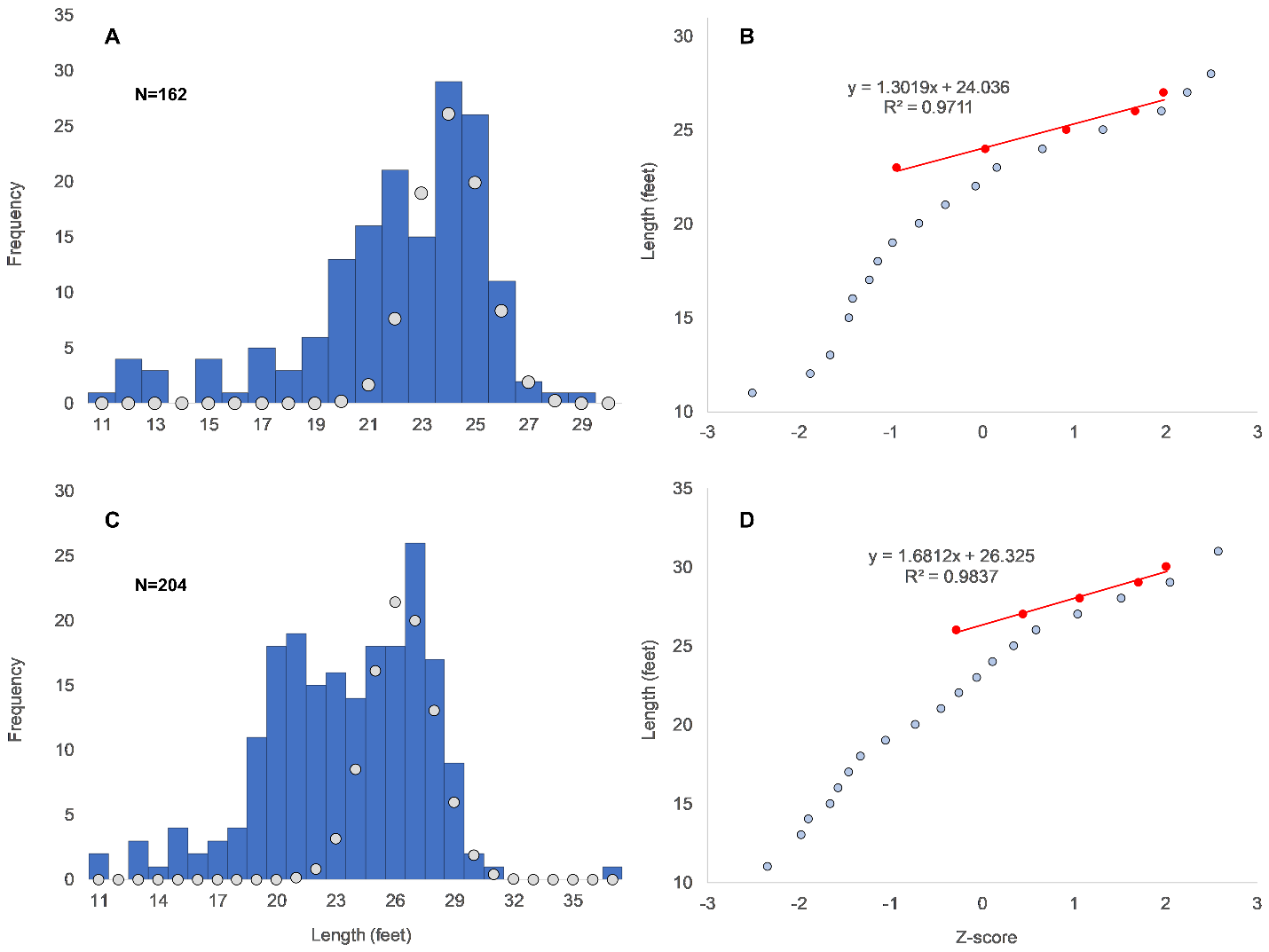

#### **Figure S4.** Length distributions and normal probability plots for female (A&B) and male (C&D) Baird’s beaked whales. Approximately 74% of males and 75% of females were mature.

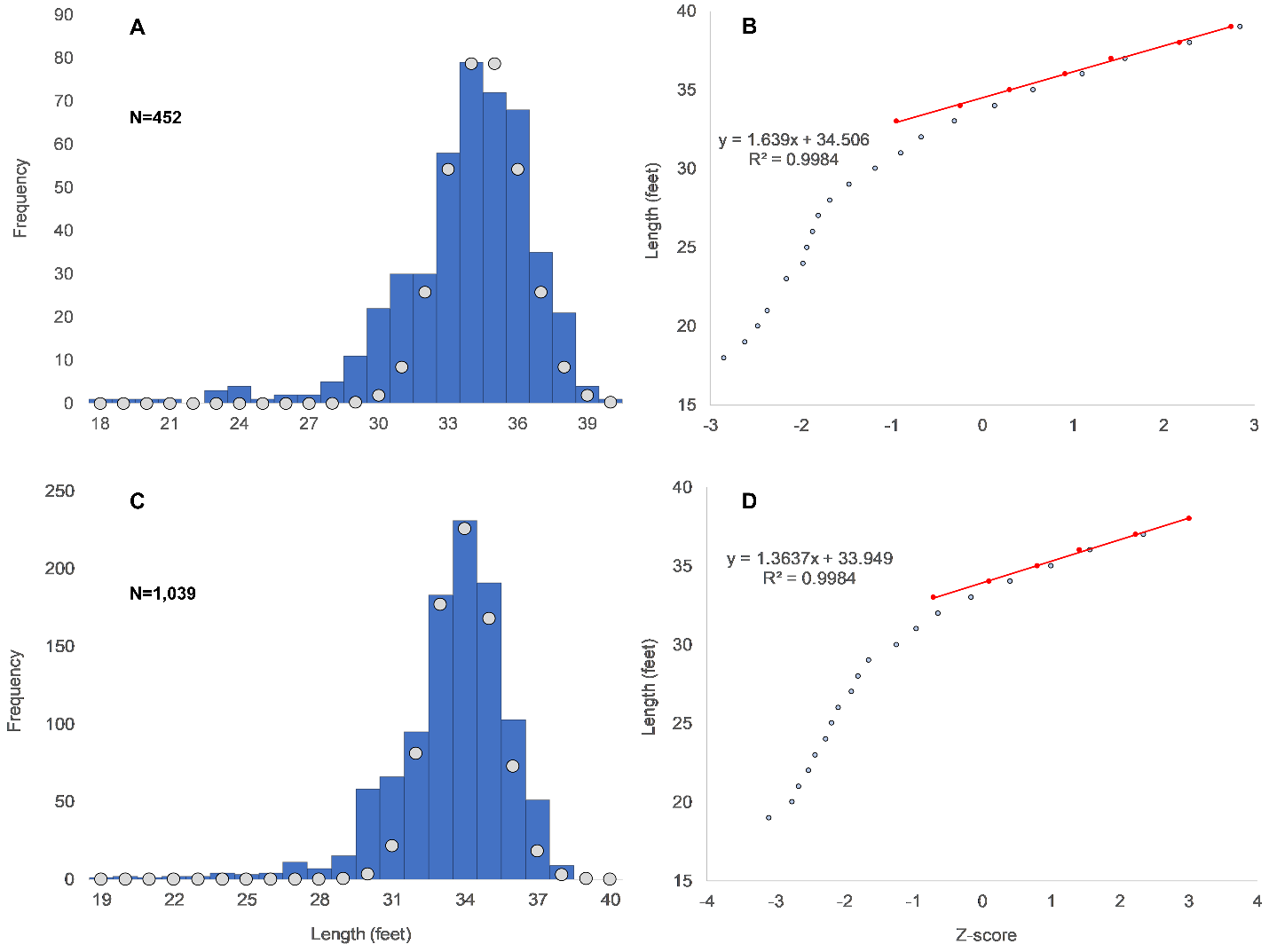

#### **Figure S5.** Linear regression of log-transformed length and mass of beaked whales (y=2.909x -1.922, n=160, r^2^=0.955). Red dotted lines are 95% prediction interval. See Supplementary File 2 for data and File 3 for references.

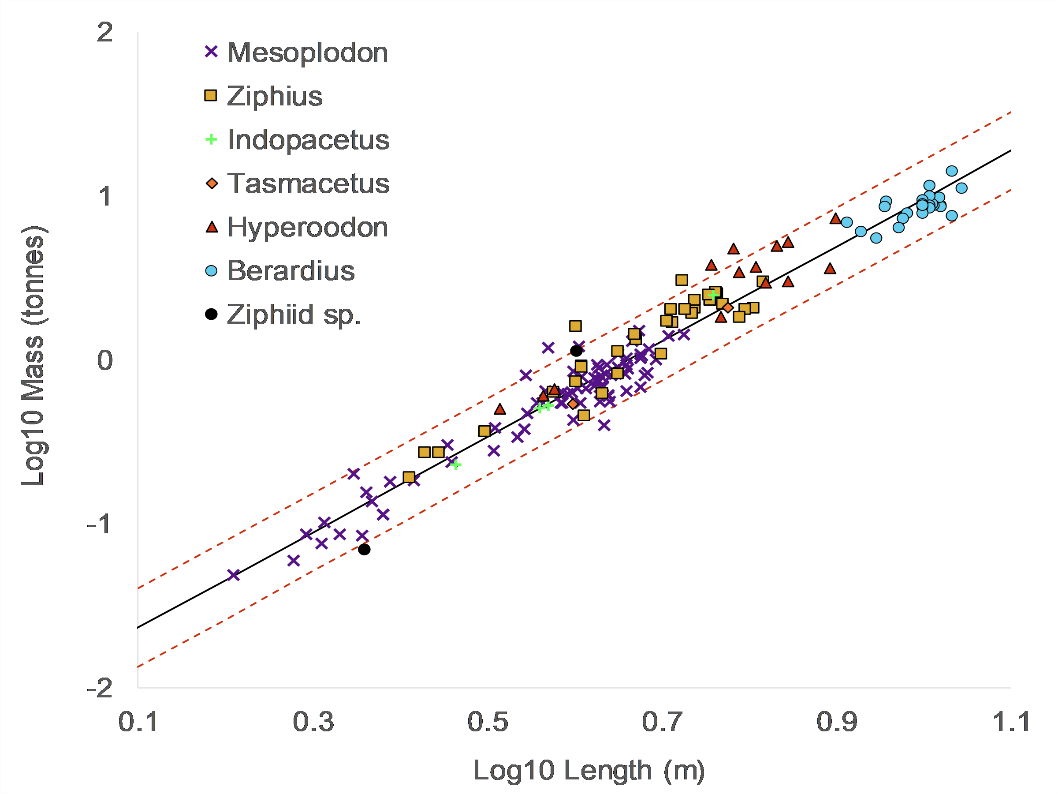

#### **Figure S6**. Normal probability plots and length distributions for global catch of mature female sperm whales (A), total female catch in the North Atlantic between 1937-71 (B&C), and global catch of males (D&E). Colored circles are fitted curves for mature whales. Approximately 45% of all reported males were mature and approximately 5% of reported females from the pre-IOS North Atlantic catches were likely misidentified males.

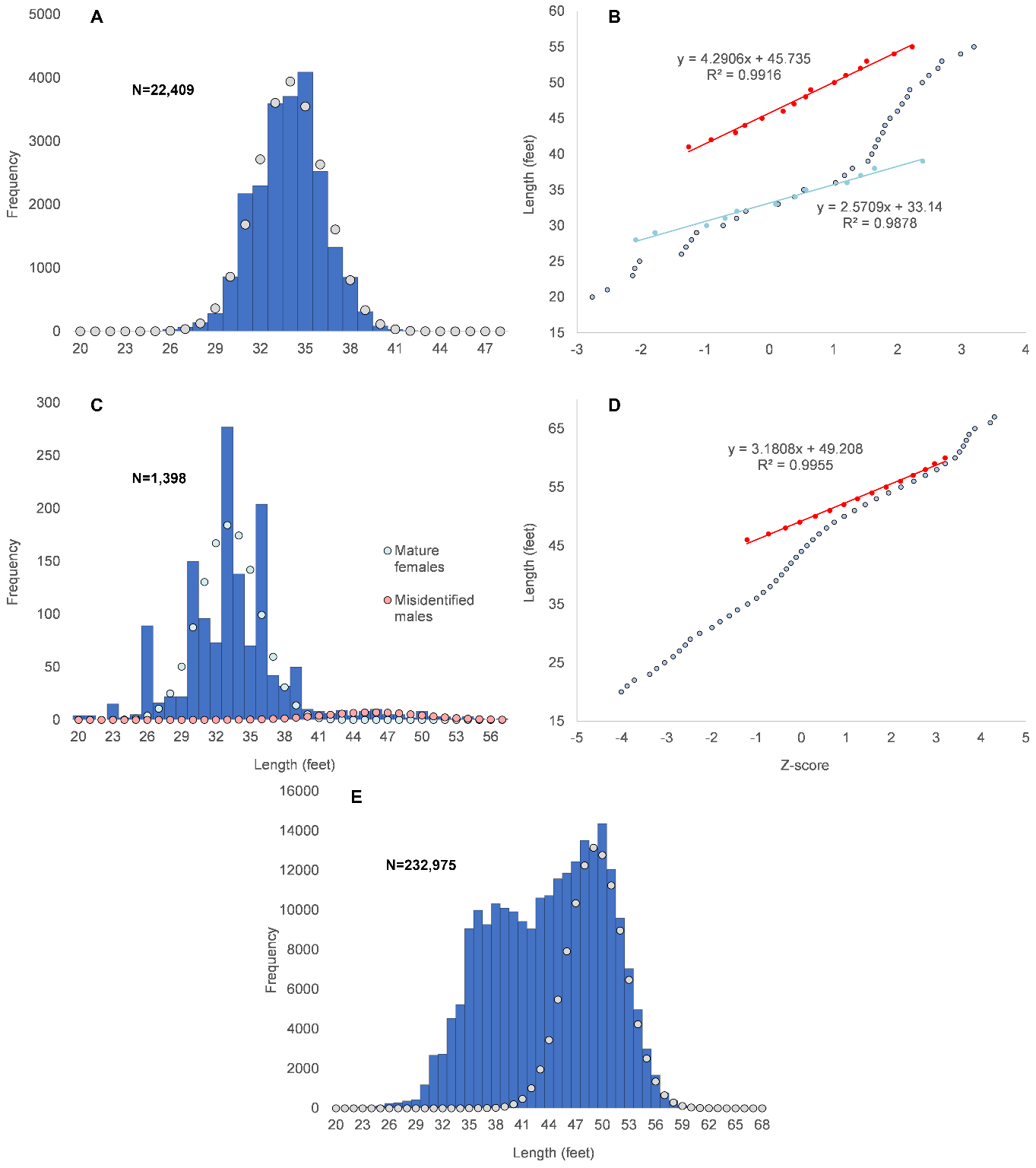

#### **Figure S7**. Relationship between the log10 total length (TL) and the following morphometric characters: (A) projection of snout (SP) in male sperm whales (y=1.629x - 1.853, n=236, r^2^= 0.547); (B) Fluke span in both sexes (y=0.883x - 0.646, n=68, r^2^= 0.846); (C) Mandible length (y=0.699x + 0.783, n=15, r^2^= 0.965); (D) Snout-angle of gape distance in males (y=0.566x + 0.854, n=222, r^2^=0.897); (E) Snout-eye distance in males (y=0.592x + 0.806, n=216, r^2^= 0.919); (F) Snout-pectoral fin tip distance in males (y=0.733x + 0.570, n=213, r^2^=0.941). (G) Maximum snout width in both sexes (y=0.518x + 1.143, n=10, r^2^= 0.967). A 17.7 m mounted skeleton from the Port Elizabeth Museum (Wood, 1972) would have a standard length of 19.2 m (80% PI: 18.9-19.7 m) based on the regression for the snout projection. The mandibles (4.9-5.5 m) and fluke spans (5.1-5.5 m) of whales allegedly measuring 24-27.4 m (Starbuck, 1878; Wood, 1982) would correspond to standard lengths of 18.4 m (80% PI: 17.3-19.7 m) to 20.0 m (80% PI: 18.7-21.4 m). See Supplementary File 2 for data and File 3 for references.

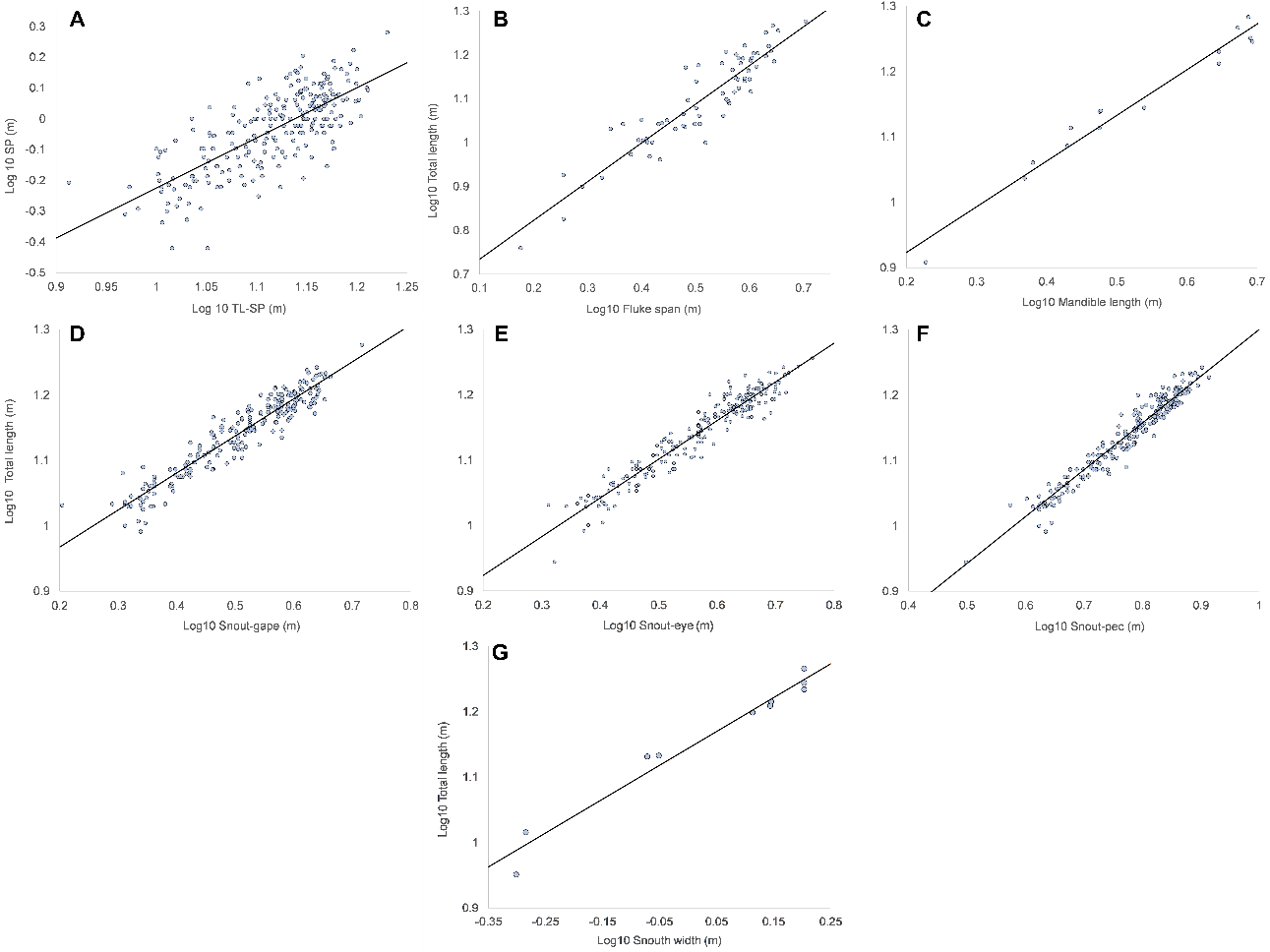

#### **Figure S8.** (A) Posterior view of the bull sperm whale stranded in Port Arthur, Texas on March, 1910 that measured 19.36 m from the snout to the end of the flukes (Newman, 1910; UNT Libraries). Fluke notch was estimated to be approximately 0.5 m. (B) Photo of the anterior body of the bull captured by the Norrøna Whaling Station in Ronas Voe, Shetland (Gordon, 1906). Measurements were taken in Aragoj (Aleixo et al, 2020) with the hat-inclusive height of the man at the center set to two scales: 175 cm and 195 cm. To account for parallax error, direct measurements other than snout width were divided by cos(θ), where θ=deviation of the head from being perpendicular to the camera. (C) Using the front of the mouth and tooth sockets as a reference, a 3D model of a sperm whale (credit to Ayla Parker) was rotated by 5° increments in Blender 3.5.1, and it was determined that θ fell between 60-65°, which was approximated to 62.5°. Results are listed in Table S1.

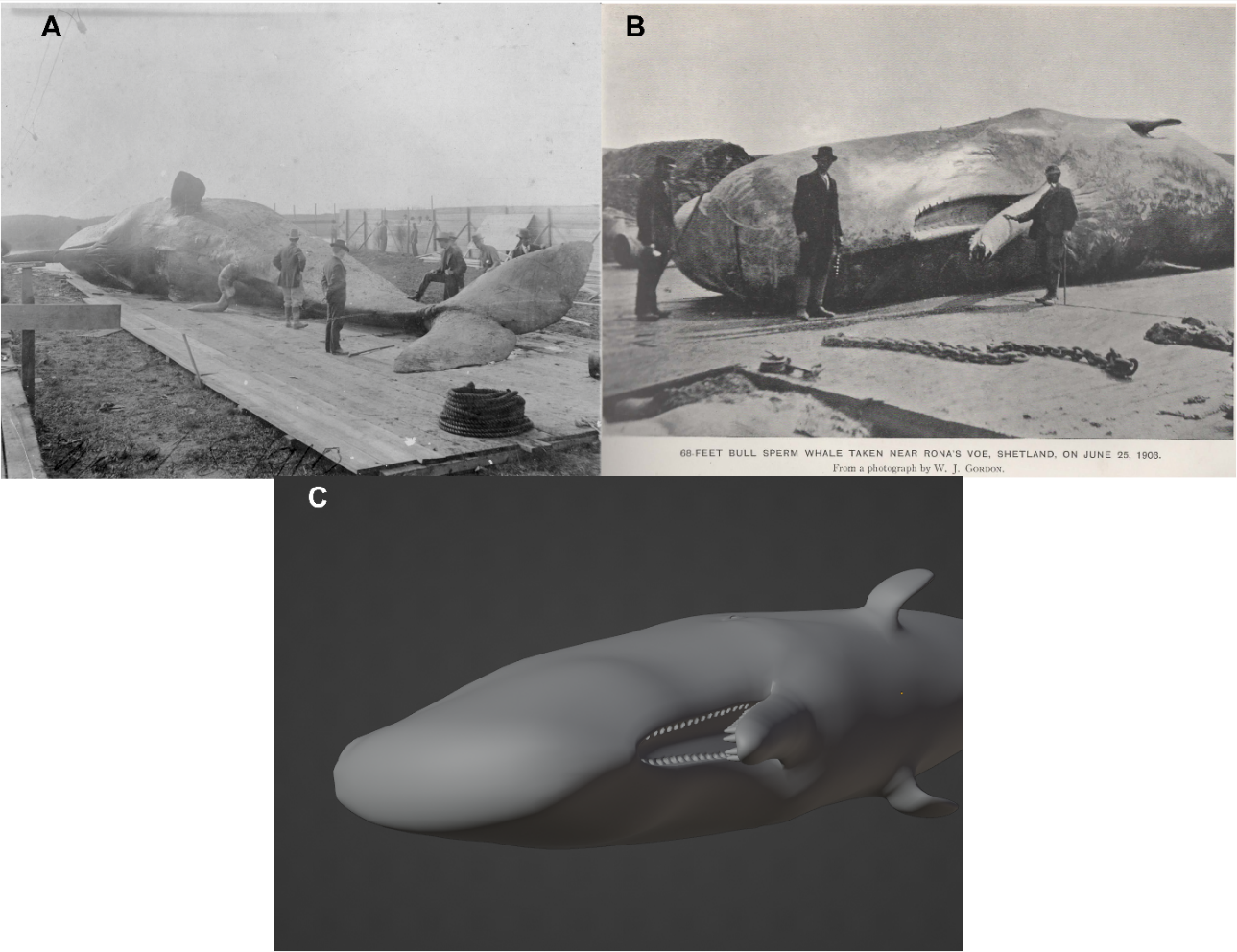

#### Table S1. Measurements and estimated TLs for Shetland bull

| Measurement | Distance (m)  Scale: 175 cm / 195 cm | Est. TL (80% PI) (m)  Scale: 175 cm | Est. TL (80% PI) (m)  Scale: 195 cm |
| --- | --- | --- | --- |
| Snout-gape | 5.6 / 6.2 | 19.0 (17.2-20.9) | 20.1 (18.2-22.1) |
| Snout-eye | 6.4 / 7.2 | 19.2 (18.2-20.2) | 20.5 (19.5-21.7) |
| Snout-Pec | 9.0 / 10.0 | 18.6 (17.8-19.5) | 20.1 (19.2-21.0) |
| Snout width | 1.8 / 2.0 | 18.9 (17.5-20.3) | 19.9 (18.5-21.4) |

#### Figure S9. Linear weight-length regression from whole weight data for sperm whales (y=2.9109x-1.8538, n=54, r^2^= 0.992). See Supplementary File 2 for data and File 3 for references.

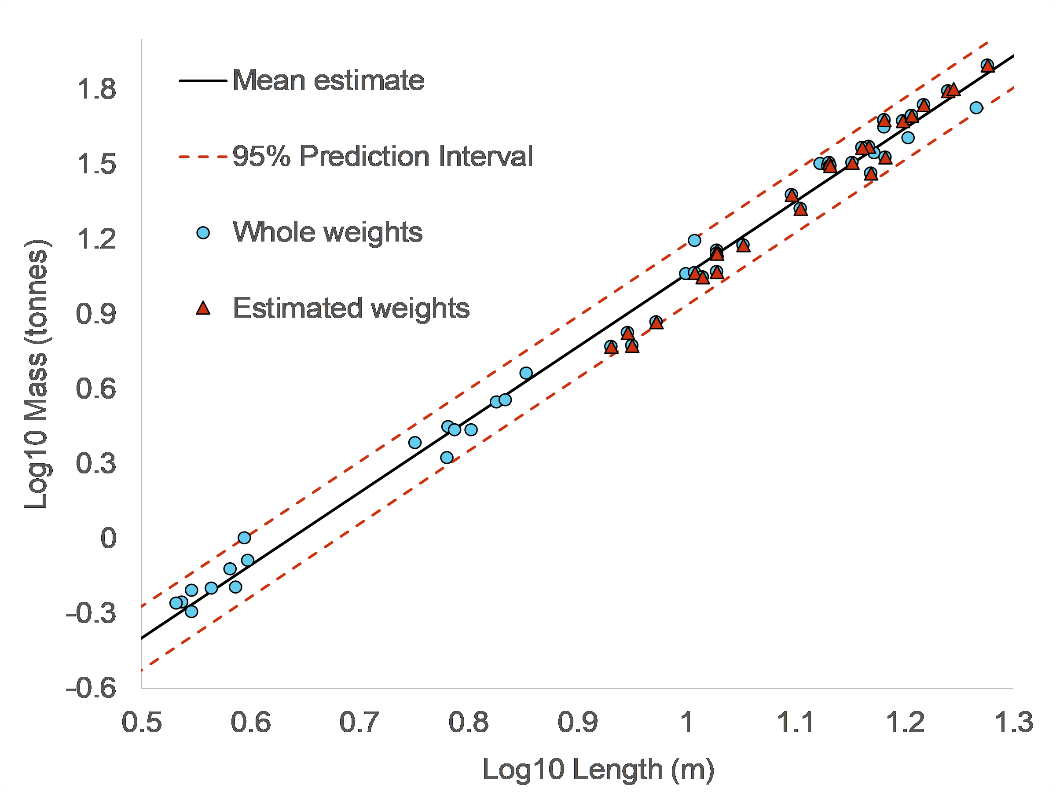
