## Supplementary File 3 for "A summary of intraspecific size variation for large odontocetes"

### **A summary of intraspecific body size variation of large odontocetes-Supplementary File 3**

Authors: Joseph McClure*

*Correspondence

564 East McIver Road, Florence, South Carolina 29506
