## Supplementary File 4 for "A summary of intraspecific size variation for large odontocetes": Supplementary File 4.R

```
##############################Orca weight analysis

###Import orca weight spreadsheet after renaming to ''orcaweight.csv)''

model1 <- lm(log10W ~ log10L, data = orcaweight)
model2 <- lm(log10W ~ log10L + Ecotype, data = orcaweight)

library(AICcmodavg)

models <- list(model1, model2)

mod.names <- c('Ecotypes pooled', 'Separate ecotypes')

aictab(cand.set = models, modnames = mod.names)

##############################Ziphiid weight analysis

###Import ziphiid weight spreadsheet, after renaming to ''ziphiidmass.csv)''

model1 <- lm(log10W ~ log10L, data = ziphiidmass)
model2 <- lm(log10W ~ log10L + Species, data = ziphiidmass)

library(AICcmodavg)

models <- list(model1, model2)

mod.names <- c('Pooled species', 'Separate species')

aictab(cand.set = models, modnames = mod.names)

########run again using ziphiidmass2
ziphiidmass2<-subset(ziphiidmass, Species!= "B.berardius")


##############################Sperm whale weight analysis

###Import sperm whale weight spreadsheet after removing whales <10 m and 
####renaming to ''spermmass.csv)''


model1 <- lm(log10W ~ log10L, data = spermmass1)
model2 <- lm(log10W ~ log10L + Method, data = spermmass1)


library(AICcmodavg)

models <- list(model1, model2)

mod.names <- c('Pooled', 'Separate methods')

aictab(cand.set = models, modnames = mod.names)

############################Sperm whale vs Ziphiid comparisons############
#####Import "Odontmass" dataframe from Supp File 2


model1 <- lm(log10W~log10L, data = Odontmass)
model2 <- lm(log10W~log10L+Clade, data = Odontmass)

library(AICcmodavg)

models <- list(model1, model2)

mod.names <- c('Clades pooled', 'Separate clades')

aictab(cand.set = models, modnames = mod.names)


########## Multiple regression with dummy variable for clade (Ziphiids vs Physeter)###########

Odontfit<-lm(log10W~log10L+Clade+log10L*Clade, data=Odontmass)
summary(Odontfit)
```
